# SlARF10, an auxin response factor, is required for chlorophyll and sugar accumulation during tomato fruit development

**DOI:** 10.1101/253237

**Authors:** Lihua Mei, Yujin Yuan, Mengbo Wu, Zehao Gong, Qian Zhang, Fengqing Yang, Qiang Zhang, Yingqing Luo, Xin Xu, Wenfa Zhang, Mingjun Miao, Zhengguo Li, Wei Deng

## Abstract

Tomato green fruits photosynthesis contributes to fruit growth and carbon economy. Tomato auxin response factor 10 (SlARF10) is one of the members of ARF family. Our results showed that SlARF10 locates in the nucleus and has no transcriptional activity. *SlARF10* was expressed in various tomato tissues, but highly expressed in green fruit. Up-regulation of *SlARF10* produced dark green phenotype of fruits, whereas down-regulation of *SlARF10* had light green phenotype. Autofluorescence and chlorophyll content analysis confirmed the phenotypes, which indicated that *SlARF10* plays an important role in chlorophyll accumulation in tomato fruits. Up-regulation of *SlARF10* increased the photochemical potential in tomato leaves and fruits. Furthermore, the *SlARF10* up-regulating lines displayed improved accumulation of starch in fruits, whereas *SlARF10* suppressed lines had inhibited starch accumulation. Up-regulation of *SlARF10* increased the expression of *AGPases*, the starch biosynthesis genes. *SlARF10* up-regulating lines had increased accumulation of *SlGLK1* and *SlGLK2* transcripts in fruits. The promoter sequence of SlGLK1 gene had two conserved ARF binding sites. *SlARF10* may regulate the expression of *SlGLK1*, thus controlling chlorophyll accumulation, photosynthesis rates and sugars synthesis in fruits. Our study provided more insight on the link between auxin signaling, chloroplastic activity and sugar metabolism during the development of tomato fruits.

**Abbreviations:**

ARFs: Auxin Response Factors
RNAi: RNA interference
GLK: GOLDEN2-LIKE
DET1/hp2: The DE-ETIOLATED 1
DDB1: UV-DAMAGED DNA-BINDING PROTEIN 1
KNOXs: Class I KNOTTED1-LIKE HOMEOBOX
GC-MS: Gas Chromatography-Mass Spectrometry
qRT-PCR: Quantitative real time PCR
TFs: Transcription factors
WT: Wild-type
MR: Middle region
DB domain: DNA binding domain
CTD: C-terminal interaction domain
AD: Transcriptional activators
RD: Transcriptional repressors
B3: N-terminal DNA-binding domain

**Highlight:** SlARF10 played an important role in the chlorophyll accumulation and photosynthesis in tomato fruits. SlARF10 was involved in starch accumulation by controlling the expression of starch synthesis related enzyme genes. *SlARF10* may regulate the expression of *SlGLK1*, thus controlling chlorophyll accumulation, photosynthesis rates and sugars synthesis in tomato fruits.

## Introduction

Tomato (*Solanum lycopersicum*) a multicarpellar berry with strong adaptability, high yield, nutrient-rich, widely used, has become the world’s second largest vegetable crop (Tanksley, 2004). Tomato fruit has arisen as the research model species for fleshy fruits, due to a short life cycle, self-pollination, and ease of mechanical crossing and genetic transformation (Klee and Giovannoni, 2011).

Fruit development can be divided into three main stages (Ho and Hewitt, 1986). The first stage is characterized by an intense mitotic activity, with an increased cell number and starch accumulation (Ho, 1996). Cell enlargement associated with the degradation of starch into soluble sugars, is characterized at the second stage of fruits (Schaffer and Petreikov, 1997). The third stage corresponds to the fruit ripening, associated with the conversion from chloroplast to chromoplast and accumulation of carotenoids, sugars, organic acids, and volatile aroma compounds in the fruit cells (Klee and Giovannoni, 2011). The accumulation of soluble solids in ripening tomato fruit is related to the starch level in immature and mature green fruit (Davies and Cocking, 1965). It was reported that between 10% and 15% of the total carbon of the fruit growth and net sugar accumulation has been contributed from photosynthetic activity in the fruit itself (Tanaka *et al*., 1974; Obiadalla-Ali *et al*., 2004). Thus chloroplast development and photosynthetic activity of green fruits affect the composition and quality of ripening tomato fruit (Nadakuduti *et al*., 2014).

It has been reported that several genes influence the development of fruit chloroplasts and the subsequent quality of ripening fruit in tomato. The DE-ETIOLATED 1 (DET1/hp2) and UV-DAMAGED DNA-BINDING PROTEIN 1 (DDB1/hp1) genes encode negative regulators of photomorphogenesis. Down-regulation of DET1/hp2 and DDB1/hp1 genes increased number of chloroplasts and plastid compartment size, leading to fruits with higher levels of chlorophyll and carotenoids in tomato fruits (Liu *et al*., 2004; Kolotilin *et al*., 2007; Rohrmann *et al*., 2011). GOLDEN2-LIKE (GLK) transcription factors are required for chloroplast and chlorophyll levels (Waters *et al*., 2008). Tomato contains two GLKs, GLK1 and GLK2, which encode functionally similar peptides. Differential expression renders *GLK1* more important in leaves and *GLK2* is predominant in fruit. The latitudinal gradient of *GLK2* expression affects the typical uneven coloration of green and ripe wild type fruit of tomato (Nguyen *et al*., 2014). Tomato ARABIDOPSIS PSEUDO RESPONSE REGULATOR 2-LIKE (SlAPRR2-like) is closest global relative of SlGLK2. Overexpression of APRR2-like gene in tomato produced larger and more numerous chloroplasts, and consequently higher chlorophyll levels in green fruits and higher carotenoid amounts in red ripening fruits (Pan *et al*., 2013). Two Class I KNOTTED1-LIKE HOMEOBOX (KNOX) proteins, TKN2 and TKN4 positively influence *SlGLK2* and *SlAPRR2-LIKE* expression to promote fruit chloroplast development in tomato fruit (Nadakuduti *et al*., 2014).

Phytohormones were reported to be involved in chloroplast development and the quality of ripening fruit (Martineau *et al*., 1994; Galpaz *et al*., 2008; Sagar *et al*., 2013). Studies of the auxin signaling transduction pathway indicated that auxin response factors (ARFs) are required for auxin-dependent transcriptional regulation in plant, and ARFs can function as either transcriptional activators or repressors of auxin-responsive genes (Ren *et al*., 2011). Most ARF proteins contain an N-terminal DNA-binding domain (B3) involved in transcription of auxin response genes, a middle region acting as an activation domain (AD) or repression domain (RD), and a C-terminal dimerization domain (Aux/IAA) requiring the formation of heterodimers or homodimers (Zouine *et al*., 2014). An increasing number of studies demonstrate that ARFs play important roles in many developmental processes of tomato (Krogan *et al*., 2011; Wang *et al*., 2012; Guan *et al*., 2013; Ckurshumov *et al*., 2014; Liu *et al*., 2014; Zhang *et al*., 2015). SlARF7 acts as a negative regulator of fruit set and development in tomato (De Jong *et al*., 2009). ARF6 and ARF8 have important roles in controlling flower growth and development (Liu *et al*., 2014). SlARF9 is required for regulation of cell division during early tomato fruit development (De Jong *et al*., 2015). SlARF3 is involved in the formation of epidermal cells and trichomes (Zhang *et al*., 2015). ARF4 was reported to control the accumulation of chlorophyll and starch in the tomato fruit (Jones *et al*., 2002; Sagar *et al*., 2013). The influence of ARF4 on fruit chlorophyll accumulation seems to be mediated through the transcriptional up-regulaton of SlGLK1 in the fruit of tomato (Sagar *et al*., 2013).

Hendelman *et al*. (2012) reported that SlARF10 is posttranscriptionally regulated by Sl-miR160, and constitutive expression of the *mSlARF10* (Sl-miR160a-resistant version) produced narrow leaflet blades, sepals and petals, and abnormally shaped fruit in tomato plants. Repression of SlARF10 expression by Sl-miR160 is essential for auxin-mediated blade outgrowth and early fruit development (Hendelman *et al*., 2012). In the present study, the functions of *SlARF10* were studied in the development of tomato fruit. Our results indicated that SlARF10 gene is involved in chlorophyll and sugar accumulation in tomato fruit. This study expand our understanding of functions of ARFs during the development of tomato fruit and provide new insight into the regulation mechanism of the chlorophyll and sugar accumulation in tomato fruit.

## Materials and methods

### Plant Materials and Growth Conditions

Tomato (*Solanum lycopersicum* L. cv. Micro-Tom) plants were grown under culture chamber conditions with 16 h light (25±2°C)/8 h dark (18±2°C) and 80% relative humidity.

### Analysis of expression patterns

The expression pattern was analyzed online using the tomato gene expression database (<u>http://gbf.toulouse.inra.fr/tomexpress/www/welcomeTomExpress.php</u>). Total RNA was extracted using a Plant RNeasy Mini kit (Qiagen). qRT-PCR was carried out as described previously (Deng *et al*., 2012).

### Subcellular localization of SlARF10

To construct SlARF10-GFP fusion expression vector, the forward 5’-ATGAAGGAGGTTTTGGAGAAGTG-3’ and reverse 5’-CTATGCAAAGATGCTAAGAGGTC-3’ primers were used to amplify the sequence of *SlARF10* coded frames. Protoplasts were obtained from suspension-cultured tobacco (*Nicotiana tabacum*) Bright Yellow-2 cells and transfected by SlARF10-GFP fusion expression vector. Transformation assays were performed as described previously (Chaabouni *et al*., 2009).

### Transcriptional activation activity of SlARF10

The ORF of *SlARF10* was amplified by using the 5’-TCCCCCGGGGATGAAGGAGGTTTTGGAGAA-3’ and 5’-CGGGATCCCTATGCAAAGATGCTAAGAGGTC-3’ primers, and fused to the GAL4 DNA-binding (DB) domain to generate pGBKT7-SlARF10 fusion construct (DB-SlARF10). The vectors were transformed into Y2H gold yeast cells and yeast cells were grown on plates with minimal medium without tryptophan (SD-W) or without tryptophan, histidine, and adenine (SD-W/H/A). The transcriptional activation activity was verified according to the growth status and activity of α-galactosidase (α-gal).

### Generation of transgenic plants

The ORF sequence of *SlARF10* was amplified by the forward 5’-TCCCCCGGGGATGAAGGAGGTTTTGGAGAA-3’ and reverse 5’-CGGGATCCCTATGCAAAGATGCTAAGAGGTC-3’ primers. The sequence was cloned into plant binary vector pLP100, resulting in overexpression vector. For construction of the RNAi vector, the 200 bp sequences of SlARF10 were amplified and the PCR products were inserted around a spacer of the β-glucuronidase gene in pCAMIBA2301 driven by a Cauliflower mosaic virus (CaMV) 35S promoter. Transgenic plants were generated via Agrobacterium tumefaciens-mediated transformation according to the method described by Jones *et al*. (2002). All experiments were performed using homozygous lines of T3 generations. For analysis of expression levels of SlARF10 in RNAi and overexpression transgenic lines, Total RNA was extracted using a Plant RNeasy Mini kit (Qiagen) and qRT–PCR was carried out as described previously (Deng *et al*., 2012).

### Analysis of chlorophyll in tomato

The chlorophyll content was measured from fruit pericarp and leaves according to the methods described by Powell *et al*., (2012). For determination of autofluorescence of chlorophylls of tomato fruits, the pericarp was peeled off tomato fruits and observed under the laser confocal microscope.

### Determination of photosynthetic substance

One gram of tomato fruits was ground by liquid nitrogen and extracted with 10ml 80% ethanol at 80℃ for 30min. After centrifuge, the super natant was dried in vacuum, evaporated to dryness and dissolved with 3 mL distilled water. One mL of dissolved samples was used for measurement of the contents of glucose, fructose, sucrose and lactose by using HPLC. The pellet of tomato fruits was used for starch analysis. Four mL of 0.2 M KOH were added to the pellet at 100℃ for 30 min. Each sample was added to 1.48 mL of 1 M acetic acid, adjusted to pH 4.5, hydrolyzed with 7 Units of amyloglucosidase for 45 min, and dissolved with 10 mL distilled water. One mL of dissolved samples was used for measurement of the starch content by using HPLC.

HPLC analysis was performed on an Agilent 1260 Series liquid chromatography system (Agilent Technologies, California, USA), which equipped with a waters XBridge Amide column (4.6×150 mm i. d., 3.5 μm) and a pre-column (Waters XBridge BEH Amide column, 3.9×5 mm i. d., 3.5 μm).

## Results

### SlARF10 belongs to ARF family, expressed mainly in tomato fruits

Amino acids sequences analysis was conducted to detect the domains of SlARF10. It was found that SlARF10 had the B3-DNA, the ARF and the AUX/IAA domains, which indicates that SlARF10 has the typical ARF conserved domains and belongs to ARF family.

The expression profiles of *SlARF10* gene in tomato plants were analyzed by online database and qRT-PCR. The database analysis revealed that SlARF10 gene is expressed in all tissues tested, including roots, stems, leaves, flowers and fruits. The expression level of SlARF10 gene is high in the fruit, especially in immature green, mature green and breaker fruits (Fig. 1A). qRT-PCR analysis also showed the similar expression profiles with high expression level of *SlARF10* in immature green, mature green and breaker fruits (Fig. 1B). The results indicate SlARF10 gene may be involved in the development of tomato fruit.

**Fig. 1.**
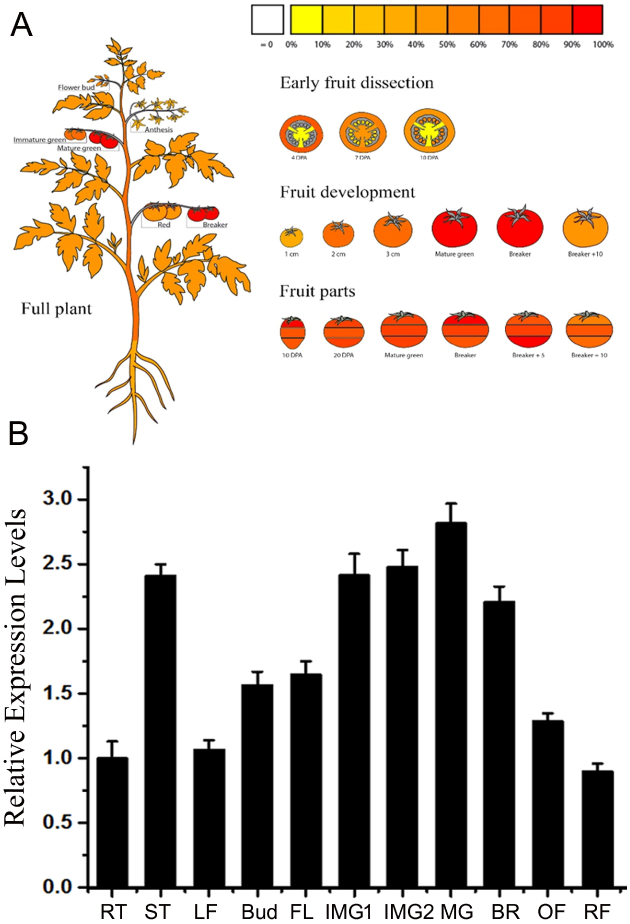
Expression pattern of SlARF10 gene in tomato plants. A, Online analysis of SlARF10 gene in tomato plants (<u>http://gbf.toulouse.inra.fr/tomexpress/www/welcomeTomExpress.php</u>). The depth of red color indicates the expression level of the gene. B, qRT-PCR analysis of expression level of *SlARF10*. The tomato housekeeping gene ubiquitin gene was used as reference. The data represent mean ±SD of three replicates.

### Subcellular localization and transcriptional activity of SlARF10

The amino acid sequence analysis found that SlARF10 has a nuclear localization signal peptide. In order to verify the location of SlARF10 in nucleus, SlARF10-GFP fusion protein vectors were constructed and transferred into tobacco protoplasts to analyze the subcellular localization of SlARF10. The green fluorescence of the SlARF10-GFP fusion protein was distributed in the nucleus (Fig. 2A), which indicated that SlARF10 is located in the nucleus.

**Fig. 2.**
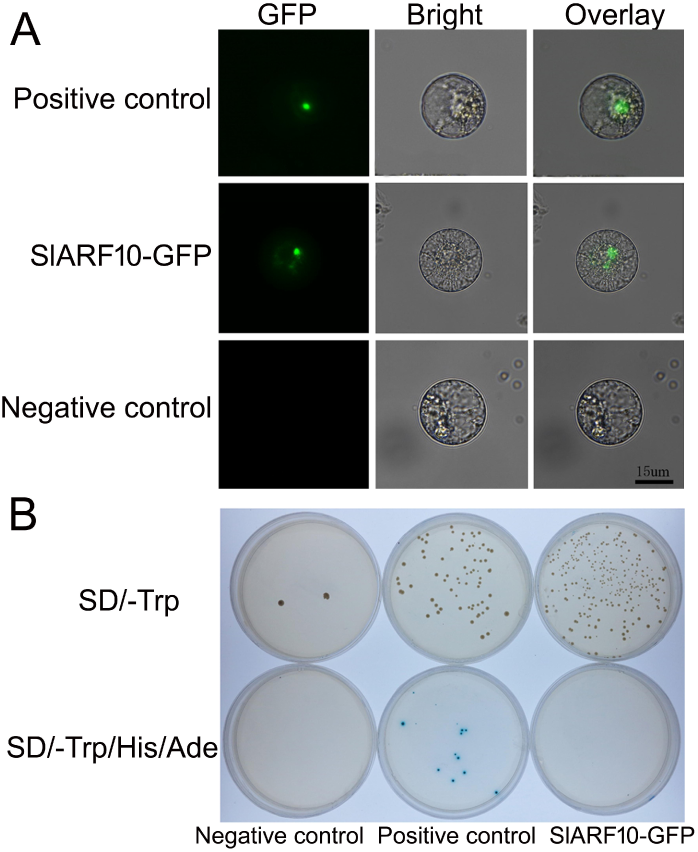
Transcriptional activation activity and subcellular localization analysis of SlARF10. A, subcellular localization analysis. PCX-DG-GFP was negative control; PCX-DG-SlARF6-GFP was positive control. Bar is 15μm. B, Transcriptional activation activity. The yeast cells, with the negative control plasmid pGBKT7, positive control pGBKT7-SlARF6 and pGBKT7-SlARF10 (right), were grown on plates with SD/-Trp or SD/-Trp-His-Ade medium.

A GAL4-repsponsive reporter system in yeast was used to analyze the transcriptional activity of SlARF10. The pGBKT7 plasmid contains the DNA binding domain (BD domain) and SlARF10 was fused to the GAL4-BD to generate pGBKT7-SlARF10 fusion plasmid and transformed into yeast. As shown in Fig 2B, the transformed yeast cell containing pGBKT7-SlARF10 recombinant plasmid could not grow on the medium lacking Trp, His, and Ade (SD-W/H/A), which is same with the yeast cell harbouring pGBKT7 plasmid (negative control). This result indicated that SlARF10 may be a transcriptional repressor.

### SlARF10 is involved in chlorophyll accumulation in tomato fruits

In order to elucidate the functions of SlARF10 gene in the development of tomato fruit, up-regulation and down-regulation of SlARF10 in tomato plants were obtained by using transgenic techniques. Ten homozygous down-regulated transgenic lines (RNAi-SlARF10) and eleven homozygous up-regulated lines (OE-SlARF10) were generated corresponding to independent transformation events. The T2 RNAi-SlARF10 and OE-SlARF10 transgenic lines with lower and higher accumulation of *SlARF10* transcripts, respectively, were selected for further study (Fig. 3A). The OE-SlARF10 lines had a dark-green fruits, while the RNAi-SlARF10 lines had light-green fruits compared with wild-type (WT) plants at green fruit stage (Fig. 3B). Moreover, the fruit colors of the transgenic lines were not significantly different with the WT lines at breaker, orange and red ripe stages (Fig. 3B).

**Fig. 3.**
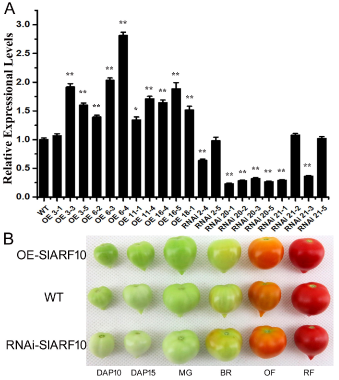
Generation of *SlARF10* transgenic plants and fruit phenotypes. A, qRT-PCR analysis of the expression of *SlARF10* in transgenic lines. B, fruit phenotypes. WT, wild type plants, OE-SlARF10, SlARF10 overexpression lines, RNAi-SlARF10, SlARF10 RNAi lines. DAP, days after pollination. MG, mature green fruit; BR, breaker fruit; OF, orange fruit; R, red fruit. The data represent mean ±SD of three replicates. ^“*”^ and ^“**”^, significant difference between transgenic and WT plants with P <0.05 and P <0.01, respectively, as determined by t-test.

Furthermore, the chlorophyll contents of green fruit and leaves were analyzed in SlARF10 transgenic plants. The RNAi-SlARF10 and OE-SlARF10 transgenic lines showed obviously lower and higher accumulation of chlorophyll content, respectively, in green fruit and leaves (Fig. 4A, 4B). Moreover, confocal laser scanning microscopy was used to detect the autofluorescence of chlorophylls in pericarp of tomato fruits. The OE-SlARF10 lines had strong chlorophylls autofluorescence, whereas the RNAi-SlARF10 lines had week autofluorescence in pericarp of green fruits (Fig. 4C). Our results indicated that *SlARF10* is involved in the chlorophyll accumulation and regulation of *SlARF10* can control the chlorophylls contents in tomato fruit.

**Fig. 4.**
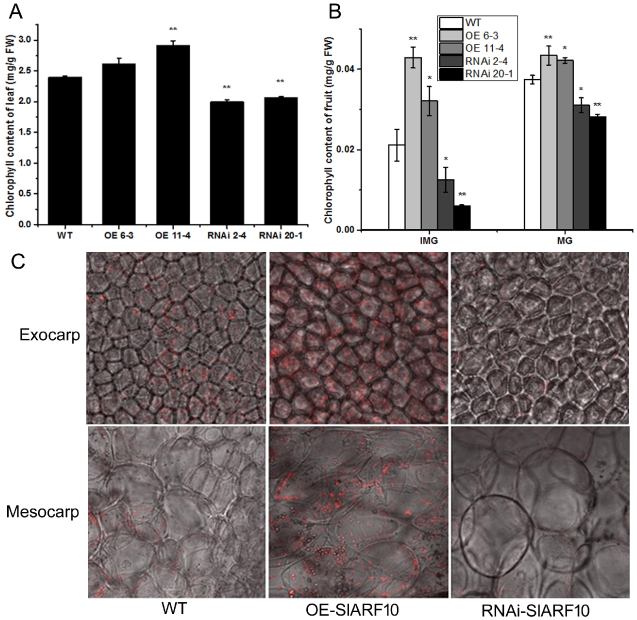
Chlorophyll accumulation in SlARF10 transgenic plants. A-B, chlorophyll contents in leaves and fruits of OE-SlARF10 and RNAi-SlARF10 plants. The data represent mean ±SD of three replicates. ^“*”^ and ^“**”^, significant difference between transgenic and WT plants with P <0.05 and P <0.01, respectively, as determined by t-test. C, Autofluorescence of chlorophylls in pericarp of tomato fruits determined by confocal laser scanning microscopy. OE-SlARF10, SlARF10 overexpression lines, RNAi-SlARF10, SlARF10 RNAi lines.

The increased chlorophyll content in the fruits and leaves may potentially confer higher photosynthetic performance in the transgenic plants. The photochemical potential was measured in the fruits and leaves of RNAi-SlARF10 and OE-SlARF10 lines. The OE-SlARF10 lines had increased photochemical potential in the fruits and leaves, whereas the RNAi-SlARF10 lines had decreased in leaves (Fig. 5). Totally, our results indicate that regulation of expression of SlARF10 gene can control the chlorophyll formation and photosynthesis in tomato plants.

**Fig. 5.**
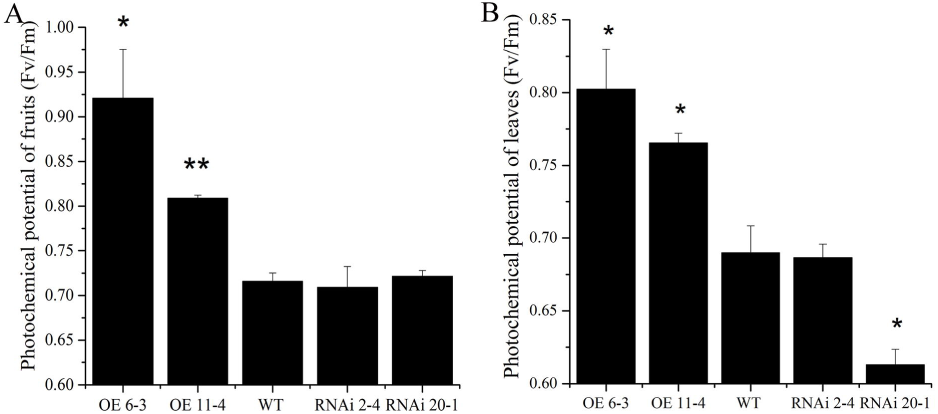
Photochemical potential in *SlARF10* transgenic plants. A, photochemical potential in fruits. B. photochemical potential in leaves. The data represent mean ±SD of three replicates. ^“*”^ and ^“**”^, significant difference between transgenic and WT plants with P <0.05 and P <0.01, respectively, as determined by t-test.

### SlARF10 affects the synthesis of photosynthetic substances in tomato fruits

Because sugar is the main product of chloroplast activity and photosynthesis, the sugar accumulation was determined in the SlARF10 transgenic plants. The cut fruits at different stages were stained with iodine to determine starch contents. The blue-purple color, indicative of the presence of starch, was mainly found in immature green fruit and mature green fruit (Fig. 6A). The OE-SlARF10 lines displayed more intense staining than that of WT plants, while the RNAi-SlARF10 showed less intense staining in green fruits (Fig. 6A). Furthermore, the starch content was measured by using HPLC method. The starch accumulated over the early green stages and rapidly degraded at the orange stage during tomato fruit development. Up-regulation of *SlARF10* obviously improved the accumulation of starch at green and breaker stages compared with WT plants (Fig. 6B), whereas down-regulation of *SlARF10* inhibited the starch accumulation at immature green stages of tomato fruits (Fig. 6C). Our results indicated that regulation of expression of SlARF10 gene controls starch synthesis in tomato green fruits.

**Fig. 6.**
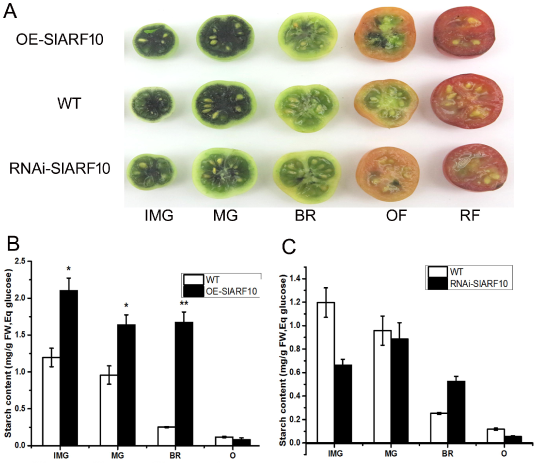
Starch accumulation in fruits of *SlARF10* transgenic plants. A, Iodine staining of tomato fruit at different developmental stages. B, starch content in transgenic plants. The data represent mean ±SD of three replicates. ^“*”^ and ^“**”^, significant difference between transgenic and WT plants with P <0.05 and P <0.01, respectively, as determined by t-test.

It is known that starch degradation is the main source of soluble sugars. We assessed the impact of up-regulation and down-regulation of *SlARF10* on the contents of fructose, glucose, sucrose and lactose in tomato fruits. OE-SlARF10 lines had significantly higher fructose content than that in the WT plants at the breaker and orange stages (Fig. 7A), whereas RNAi-SlARF10 lines had no obvious difference during tomato fruits development (data not shown). Also there were no distinct differences between WT, OE-SlARF10 lines (Fig. 7B) and RNAi-SlARF10 lines (data not shown) in glucose content. For the disaccharide, the contents of sucrose and lactose in OE-SlARF10 line were significantly higher than that in WT lines (Fig. 7C, D). In RNAi-SlARF10 lines, the two disaccharides contents were lower than WT lines during tomato fruit development (Fig. 7E, F).

**Fig. 7.**
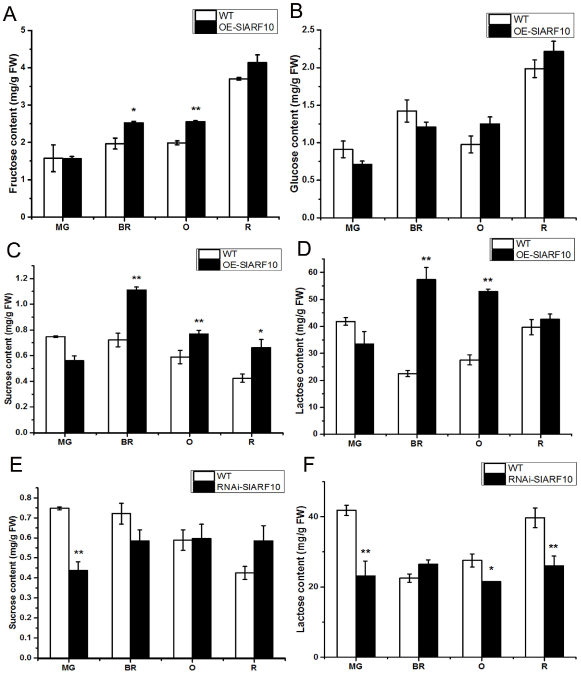
Accumulation of photosynthetic substances in fruits of *SlARF10* transgenic plants. Fructose (A) and glucose (B) contents in overexpression transgenic plants. Sucrose (C) and lactose (D) contents in overexpression transgenic plants. Sucrose (E) and lactose (F) contents in RNAi transgenic plants. The data represent mean ±SD of three replicates. ^“*”^ and ^“**”^, significant difference between transgenic and WT plants with P <0.05 and P <0.01, respectively, as determined by t-test.

### SlARF10 regulates the expression of Starch Biosynthesis Genes

To gain more insight into the mechanism of sugar metabolism in SlARF10 transgenic plants, we analyzed the expression pattern of starch biosynthesis genes. AGPase genes, with four subtypes (*AGPase-L1*, *AGPase-L2*, *AGPase-L3* and *AGPase-S1*), are the most important enzyme in starch synthesis process, which catalyzes the first step reaction of starch synthesis. *AGPase-L1*, *AGPase-L2*, *AGPase-L3* and *AGPase-S1* genes show the higher levels of expression at different fruit development stages in OE-SlARF10 plants. The expression of AGPase-L2, AGPase-L3 and AGPase-S1 were significantly higher than that in WT plants in immature green fruit stage, but the expression of AGPase-L1 was not significantly different (Fig. 8A). In the mature green fruit period, AGPase-S1 had a significantly higher expression, while the other three genes had no obvious difference (Fig. 8B). In the fruit breaker period, only AGPase-L1 and AGPase-L2 genes displayed higher expression levels compared with WT plants (Fig. 8C). These results indicated that up-regulation of SlARF10 gene improve the expression of AGPase genes.

**Fig. 8.**
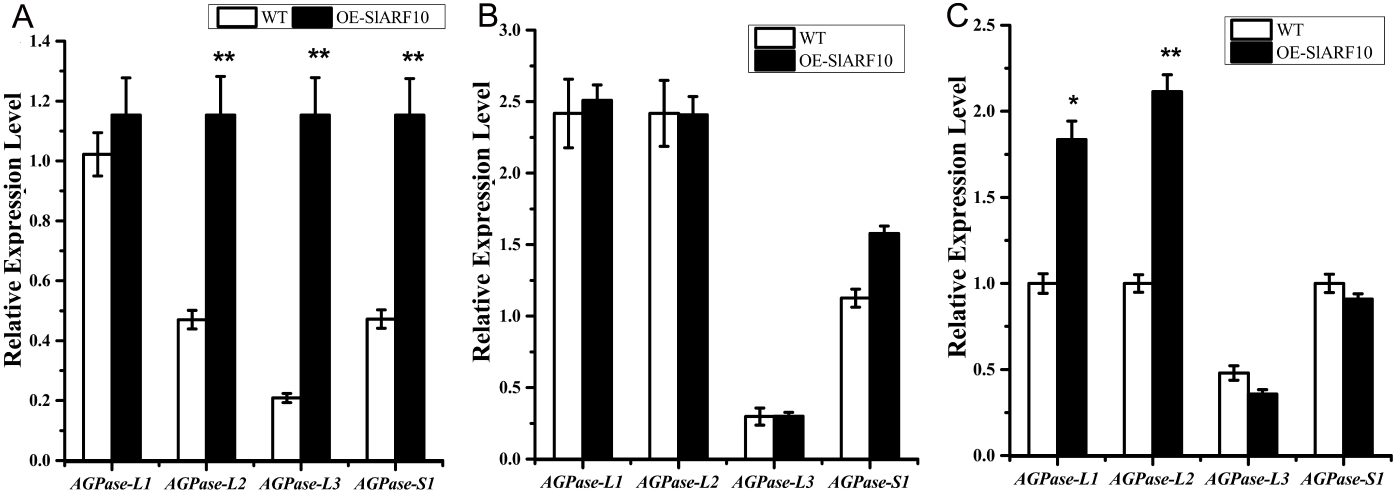
The expression of SlAGPase genes in SlARF10 transgenic plants. The levels of transcripts were assessed in tomato fruit by RT-PCR at IMG, MG, BR stage for SlAGPaseL1 (L1), SlAGPaseL2 (L2), SlAGPaseL3 (L3), and SlAGPaseS1 (S1). The data represent mean ±SD of three replicates. ^“*”^ and ^“**”^, significant difference between transgenic and WT plants with P <lt;0.05 and P <lt;0.01, respectively, as determined by t-test.

### Up-regulation of SlARF10 increased the expression levels of *SlGLK1* and *SlGLK2*

The chlorophyll and starch phenotypes of OE-SlARF10 plants are reminiscent of those described in *SlGLK* overexpression transgenic plants. The expression levels of two GLK genes, *SlGLK1* and *SlGLK2*, were analyzed in OE-SlARF10 and RNAi-SlARF10 plants. qRT-PCR showed increased accumulation of *SlGLK1* and *SlGLK2* transcripts in the fruits of OE-SlARF10 plants and decreased accumulation of the transcripts in the fruits of RNAi-SlARF10 plants (Fig. 9). Analysis of the promoter sequence of SlGLK1 gene found two conserved ARF binding sites, TGTCTC box. These results indicated SlARF10 may bind to TGTCTC box, thus regulating the expression of *SlGLK1* and controlling chlorophyll accumulation. Moreover, qRT-PCR showed there is no obvious difference between the WT and transgenic plants in the expression levels of *DDB1* and *THY5* genes (Fig. 9). This result indicated that the effect of *SlARF10* on chlorophyll accumulation acts independently of *DDB1* pathway. The expression levels of protochlorophyllide reductase gene (*PR*), chlorophyll binding protein 1 gene (*CBP1*), chlorophyll binding protein 2 gene (*CBP2*) were also analyzed in the transgenic plants. The *PR*, *CBP1*, *CBP2* had increased accumulation of transcripts in the fruits of OE-SlARF10 plants and decreased accumulation in the fruits of RNAi-SlARF10 plants.

**Fig. 9.**
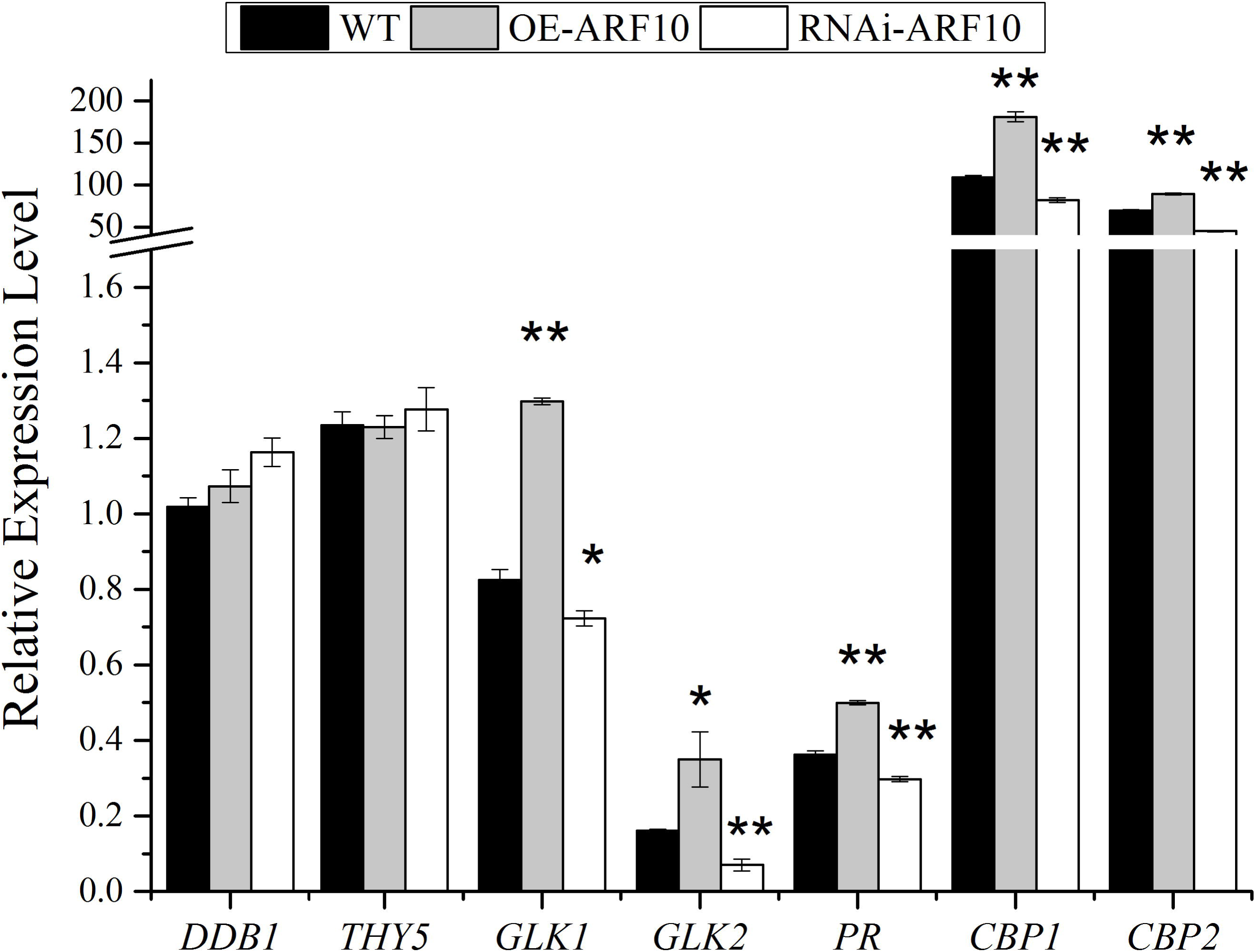
Expression profile of the genes related with chlorophyll formation in *SlARF10* transgenic tomato fruits. *DDB1*, Solyc02g021650, UV damaged DNA binding protein 1. *THY5*, Solyc08g061130, bZIP domain of plant elongated/long HY5-liketranscription factors and similar proteins gene. *GLK1*, Solyc07g053630, golden2-like protein 1 gene. *GLK2*, Solyc10g008160, golden2-like protein 2 gene. *PR*, Solyc10g006900, protochlorophyllide reductase gene. *CBP1*, Solyc02g070990, chlorophyll binding protein 1 gene. *CBP2*, Solyc02g070950, chlorophyll binding protein 2 gene. The data represent mean ±SD of three replicates. WT, Wild type plants. ^“*”^ and ^“**”^, significant difference between transgenic and WT plants with P <0.05 and P <0.01, respectively, as determined by t-test.

## Discussion

The phytohormone auxin regulates a wide variety of developmental processes by modulating gene expression via a family of transcriptional regulators, namely, Auxin Response Factors (ARFs). ARFs act as transcriptional activator or repressor of auxin-responsive genes by direct binding to the promoter (Li *et al*., 2016). Our research demonstrates that *SlARF10* scarcely has transcriptional activity. It is conceivable that ARF10 acts as a significant transcriptional repressor during plant growth and development.

Strikingly, previous studies on transactivation assays have indicated that 36% of tomato *ARFs* are strong repressors of transcriptional activity but only 22% work as transcriptional activators (Zouine *et al*., 2014). It has been reported that full-length ARF1 and ARF2 repressed transcription with or without exogenous auxin treatment in Arabidopsis (Tiwari *et al*., 2003). However, the repressor/activator ratio among *ARFs* in Arabidopsis (1.7) is less than half of that in tomato (3.6) (Zouine *et al*., 2014).

Representative ARF proteins embrace a conserved N-terminal DNA Binding Domain (DBD) that regulates the expression of early auxin response genes, a nonconserved middle region (MR) that decides whether ARFs activate or repress target genes, and in most cases a conserved C-terminal interaction domain (CTD) that contributes to mediating interactions between ARFs, as well as between ARFs and their Aux/IAA inhibitors (Guilfoyle *et al*.,2007; Boer *et al*., 2014; Kim *et al*., 1997). A preliminary conclusion based on transient expression assays can be draw that ARFs with Q-rich MRs function as transcriptional activators (AD) while a majority of other ARFs function as transcriptional repressors(RD) (Ulmasov *et al*., 1999). To gain clues on the structural feature of ARF10 function as a potential transcriptional repressor, gene structure analysis was performed to differentiate ARF10 from other activators. ARF10 harbors a predicted repression domain in the MR and hence are predicted to function as RD (Zouine *et al*., 2014), which is consistent with our speculation.

The chlorophyll content, as a critical feature of unripe fruits, affects the nutritional components and flavor of ripe fruit. Moreover, the link between chlorophyll content and photosynthesis or photosynthate metabolism in fruit tissues has been illuminated by a variety of studies (LopezJuez and Pyke, 2005; Nadakuduti *et al*., 2014; Powell *et al*., 2012), though the regulatory mechanisms by which this predominant pigment impacts photosynthetic capacity as well as photosynthate accumulation and therefore fruit quality remain unclear. Auxin plays a pivotal role in initiation of fleshy fruit development and determining final fruit size through the control of cell division as well as expansion (Sagar *et al*., 2013; Devoghalaere *et al*., 2012). Subsequently, auxin impacts an array of crucial regulators, such as ethylene, ABA and *Rin*, and vital effectors, such as genes for β-xanthophyll and lycopene biosynthesis as well as for chlorophyll degradation (Su *et al*., 2015; Manoharan *et al*., 2017). It has also been suggested that *Arabidopsis thaliana* roots, regulated by auxin treatment, demonstrate enhanced chlorophyll accumulation as well as chloroplast development after detached from shoots and then mutant analyses indicate that auxin transported from the shoot represses chlorophyll accumulation via the function of ARF7, ARF19, and IAA14 (Kobayashi *et al*., 2012). A hypothesis based on these evidences can be draw that auxin, as a critical phytohormone, regulates chlorophyll accumulation and degradation via function of ARFs during fruit setting and fruit development.

Given the experimental phenomenon that IAA14 and ARF7/19 mediate auxin signaling pathway to repress chlorophyll biosynthetic genes in Arabidopsis thaliana (Kobayashi *et al*., 2012), we speculate that auxin is likely to regulate chlorophyll biosynthesis and accumulation via activated or repressed transcriptional function of ARFs. Previous work manifested that DR12/ARF4, a member of the tomato ARF gene family of transcription factors, influences the regulation of fruit development, that is, transgenic tomato plants with down-regulated *SlARF4* expression levels bore dark-green fruit at immature stages, with significantly increased chlorophyll content, and accumulated more starch at incipient stages of fruit development as well as more sugar at the ripening stages. *SlARF4* may function through the transcriptional repression of GLK1 gene expression in tomato fruits (Sagar *et al*., 2013; Jones *et al*., 2002). Conversely, in the current research, up-regulation of *SlARF10*, another transcriptional repressor, elicits enhanced chlorophyll accumulation in tomato fruit. Also, our results showed overexpression of *SlARF10* increased accumulation of *SlGLK1* transcripts in the fruits. *SlARF10* may control chlorophyll accumulation through regulating the expression of *SlGLK1*. Our results also support the idea that transcriptional regulation of the photosynthetic activity may be through a common route in tomato fruits. It is possible that ARF10 and other ARF efficiently bind to form stable dimerization complexes, such as those found in ARF6 and ARF8 in Arabidopsis.

Chlorophyll a is initially synthesized from glutamyl-tRNAglu, and chlorophyll b is synthesized from chlorophyll at the final step of chlorophyll biosynthesis. Analysis of the complete genome of Arabidopsis thaliana elucidated that there are 15 enzymes encoded by 27 genes for chlorophyll biosynthesis (Beale *et al*., 1999; Nagata *et al*., 2013). Although the underlying mechanism for auxin controlling chlorophyll biosynthesis pathway remains poorly understood, we hypothesize that the function of ARFs, during chlorophyll biosynthesis, is likely to regulate key gene expression such as HEMA1, HEMA2, and HEMA3.The reduction of glutamyl-tRNA catalyzed by glutamyl-tRNA reductase (GluTR) which is encoded by HEMA1, HEMA2, and HEMA3, is the rate-limiting and an vital regulation step in the tetrapyrrole biosynthetic pathway (Zhao *et al*., 2014).

*SlARF10* up-regulated lines displayed dark-green fruit phenotypes in parallel with those showed by *SlARF4* down-regulated lines with enhanced chlorophyll content (Sagar *et al*., 2013). Whereas, in contrast to *SlARF4* under-expressing plants where dark-green phenotype is restricted to immature fruits, significantly higher chlorophyll content in *SlARF10* over-expressed lines was detected in both leaf and fruit tissues. This feature indicated that, in contrast with *SlARF4*, *SlARF10* control of chlorophyll accumulation is not fruit-specific. Furthermore, the higher chlorophyll content in *SlARF10* over-expressed lines correlating with a higher photochemical efficiency compared with wildtype elicits elevated starch levels and sugar content in the transgenic fruit. Although the prevailing theory is that predominant fruit growth and metabolism are sustained by photoassimilate supply from the original source (Ruan *et al*., 2012), our result cannot exclude that increased starch and sugar content in OE-SlARF10 lines could also results from a more effective transportation of photoassimilate into fruit. It is possible that enhanced leaf photosynthesis observed in up-regulated transgenic lines is a supposed supply that could provide fruit with photoassimilate. This viewpoint is consistent with experimental evidence that that down-regulation of *SlIAA9* alters auxin sensitivity and facilitates the development of vascular bundles (Wang *et al*., 2005), thereby likely increasing sink strength as well as assimilation product supply to the fruit.

Starch is not only a significant carbohydrate reserve in the majority of plant but also a predominant factor to define fruit nutrition and favor. In plant starch synthesis, the first regulatory step, the synthesis of ADP-glucose, is catalyzed by AGPase from glucose-1-phosphate and ATP (Yin *et al*., 2009; Stark *et al*., 1992). Experimental evidences were then provided showing that, in potato (*Solanum tuberosum*) tubers, this critical catalytic reaction is also the limiting step during starch biosynthesis (Tiessen *et al*., 2002). It has been reported that auxin regulates expression of the *SlAGPase* gene (Miyazawa *et al*., 1999), and indeed down-regulation of *SlARF4* increased both starch content and the expression of essential genes involved in starch biosynthesis in tomato fruit, particularly genes coding for AGPase (Sagar *et al*., 2013). In our research, the improved starch content in *SlARF10* up-regulating lines correlates well with the increased expression of *AGPase* genes in starch biosynthesis, indicating that *SlARF10* likely regulates starch accumulation via controlling *SlAGPase* gene expression. Up-regulation of *SlARF10* also leads to higher soluble sugar content at various stages of tomato fruit while down-regulation fruit displays decreased sugar accumulation, likely owing to the different content of starch which could be degraded into soluble sugars at the developmental stage of plant fruit. This is in accordance with previous studies demonstrating that incipient starch content determines soluble solid content during fruit development (Schaffer *et al*., 2000; Baxter *et al*., 2005).

Overall, the current study demonstrates that *SlARF10* gene plays a significant role in chlorophyll accumulation during fruit development in tomato. The data also has shed some light on the ability of auxin regulating starch accumulation during fruit development via altering gene expression of *SlARF10*. However, auxin regulation of carbohydrate accumulation, especially its connection with other regulatory mechanisms, are still to be elucidated. Future work will center on illuminating auxin regulatory network for chlorophyll and starch biosynthesis including reveal gene function of relevant transcriptional factors.

## Acknowledgements

This work was supported by the National Key Research and Development Program (2016YFD0400100), the Project of Chongqing Science and Technology Commission (CSTC2015JCYJA80018), the National Natural Science Foundation of China (31272165) and the National Basic Research Program of China (2013CB127106).

